# Epitope-Focused Immunogens Confer Broad Protection against Coronaviruses

**DOI:** 10.64898/2026.08.02.742375

**Authors:** Mingxi Li, Qing Chen, Junghwa Seo, Yaoming Liu, Fred Zhangzhi Peng, Kexin Huang, Ziniu Dai, Changxu Chen, Zeli Zhang, Jincun Zhao, Zhaoqian Wang, Fajie Yuan, Xiancai Ma, David R. Martinez, Dapeng Li

**Author notes:** Corresponding Authors: Xiancai Ma, David R. Martinez and Dapeng Li. These authors contributed equally to this work (Mingxi Li, Qing Chen, Junghwa Seo).

## Abstract

Variable regions in coronavirus spike dominate antibody responses elicited by infection or conventional vaccination and limit induction of broad neutralization. In contrast, conserved S2 elements, including the stem helix (SH) and fusion peptide proximal region (FPPR), are promising but subdominant targets of protective immunity. Here, we employed computational design to generate *de novo* epitope-focused immunogens that precisely present the SH and FPPR epitopes. Formulated as combinatorial immunogens, this vaccine elicited consistent serum responses with broad reactivity across known human coronaviruses and induced epitope-specific antibodies with broad neutralizing activity. As a heterologous boost, epitope-focused immunogens protected mice against challenge with phylogenetically distinct coronaviruses, including bat SARS-related RsSHC014-CoV and MERS-CoV, establishing a generalizable strategy for precision immune focusing and advancing universal coronavirus vaccine development.

## INTRODUCTION

Coronavirus spillovers and the repeated emergence of antigenically distinct variants have exposed key limitations of current spike-based vaccines. Although the existing SARS-CoV-2 vaccines protect against severe disease, their efficacy against divergent strains is constrained by antibody responses that preferentially target highly variable regions of the spike protein (*1-5*). As a result, there remains a critical need for vaccine strategies capable of redirecting humoral immunity toward structurally conserved sites of vulnerability shared across diverse coronaviruses.

Broadly neutralizing antibodies isolated from infected or vaccinated individuals have identified conserved epitopes in the spike S2 subunit, including the stem helix domain (residues 1145-1156) (*6-9*) and fusion peptide proximal region (residues 815-823) (*10-12*). These regions are conserved across multiple human and zoonotic coronaviruses. However, the stem-helix domain and fusion peptide proximal region are typically subdominant during natural infection and vaccination, suggesting that their weak immunogenicity reflects limitations in antigen presentation rather than lack of protective potential (*13, 14*). This disconnect between conserved antigenic vulnerability and immune focusing indicates that humoral responses are shaped not only by antigen sequence alone, but also by epitope accessibility and structural context within immunogens.

Here, we hypothesized that structurally defined epitope scaffolding could overcome this limitation by extracting conserved neutralizing determinants from the full spike protein and presenting them on *de novo* designed scaffold proteins in optimized conformations. We show that epitope-focused immunogens targeting conserved S2 regions can redirect vaccine-elicited immunity toward broadly protective antibody responses and confer cross-lineage protection across divergent coronaviruses such as bat SARS-like zoonotic coronaviruses and MERS-CoV.

## RESULTS

### *De novo* generation of epitope-focused immunogens

To develop broadly protective coronavirus immunogens, we focused on two conserved neutralizing epitopes in the spike S2 subunit: the stem helix (SH; residues 1146–1159) and the fusion peptide proximal region (FPPR; residues 813–825) (***Fig. 1A***). Both epitopes are targeted by previously reported broadly neutralizing antibodies (bnAbs), including S2P6, DH1057, WS6, 76E1, COV44-62, and VN01H1 (*6, 8, 10-12, 15*), identifying them as con-served sites of coronavirus vulnerability. Sequence analysis across seven human coronaviruses showed substantial conservation of FPPR across both alpha- and betacoronaviruses, whereas SH is more divergent in distantly related viruses, particularly HKU1, 229E, and NL63 (***Fig. 1A***).

**Fig. 1.**
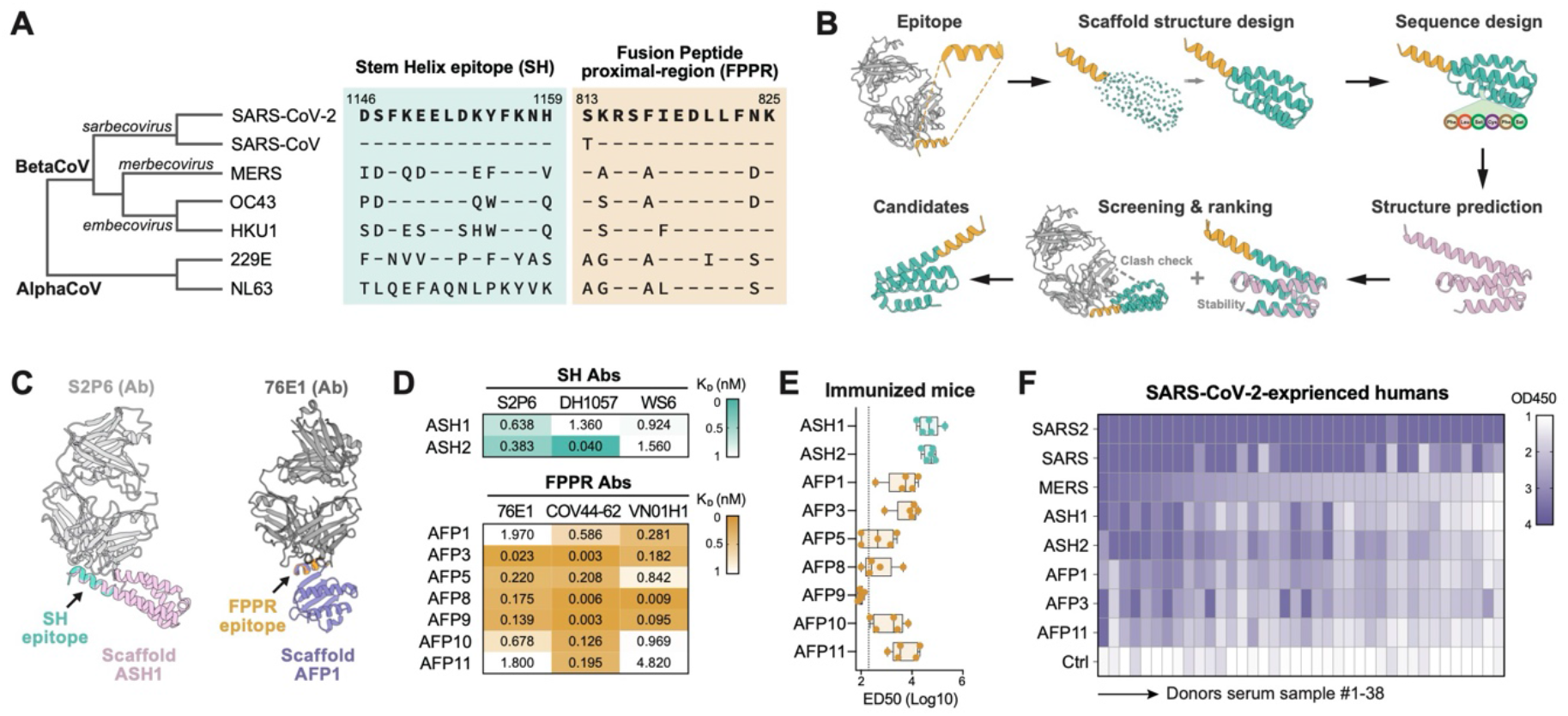
*De novo* generation of stem helix (SH) and fusion peptide-proximal region (FPPR) epitope scaffolds. **(A)** Maximum-likelihood cladogram of representative alpha- and beta-coronavirus spike protein sequences (left), with matched alignments of the SH and FPPR from each virus (right). Residues are numbered according to the SARS-CoV-2 spike epitope sequences used throughout this study. **(B)** Schematic of the *de novo* epitope-scaffold design workflow. Scaffold backbones and sequences were generated around the target epitope conformations, refolded in silico, computationally scored, and top-ranked candidates were selected for experimental screening. **(C)** Computational models of selected antibodies bound to representative SH (left) and FPPR (right) epitope scaffolds. Epitopes are shown in green (SH) and orange (FPPR). **(D)** Conformational validation by monoclonal antibody (mAb) binding. Binding affinities of SH- and FPPR-directed mAb to designed scaffolds measured by SPR and reported as K_D_ values. **(E)** Functional validation by immunization. BALB/c mice were immunized four times with designed epitope scaffolds (n = 5 per group), and serum binding to SARS-CoV-2 spike was measured by ELISA and reported as ED50 values. **(F)** Reactivity of epitope scaffolds and spike proteins with plasma from SARS-CoV-2-experienced individuals, measured by ELISA and reported as OD450 values at a 1:100 plasma dilution. Plasma samples are ordered from left to right by decreasing reactivity to MERS-CoV spike. Data are means ± SEM.

To generate SH and FPPR epitope-focused immunogens, we used a protein design pipeline to generate *de novo* scaffold immunogens that recapitulate the native conformations of SH and FPPR epitopes (***Fig. 1B***). Epitope backbone geometries derived from bnAb-bound structures were grafted onto *de novo* scaffold architectures using RFdiffusion (*16*), followed by sequence design with ProteinMPNN (*17*) and *in silico* screening with AlphaFold3 and other tools for structural stability (*18, 19*), compatibility with bnAbs (*20*), and developability properties. Representative computational models of the resulting immunogens, designated ASH for SH scaffolds and AFP for FPPR scaffolds, predicted high-fidelity structural mimicry of the native epitopes in complex with their corresponding bNAbs S2P6 and 76E1 (***Fig. 1C and Fig. S1A***). Surface plasmon resonance (SPR) confirmed that these designed ASH and AFP immunogens bound multiple SH- or FPPR-directed bnAbs with high affinity, predominantly in the sub-nanomolar to low-nanomolar range (***Fig. 1D and Fig. S1B***), indicating faithful epitope presentation and display. Immunization of BALB/c mice with monomeric ASH and AFP immunogens identified ASH1, ASH2, AFP1, AFP3, and AFP11 as the most immunogenic candidates, based on serum reactivity to SARS-CoV-2 spike (***Fig. 1E and Fig. S1C***). Circular dichroism spectra of the purified epitope scaffolds were consistent with the design models, and the melting temperatures for most were greater than 95°C (***Fig. S1D***).

We next asked whether the designed immunogens captured antigenic features associated with cross-reactive coronavirus immunity. Plasma samples from 38 individuals with prior SARS-CoV-2 vaccination and/or infection showed that strong cross-reactive binding to MERS-CoV spike also correlated with enhanced recognition of the scaffold immunogens (***Fig. 1F***). Thus, the ASH and AFP immunogens present epitopes targeted by naturally elicited broad coronavirus responses. Together, these results demonstrated that AI-guided epitope scaffolding can generate structurally precise and antigenically faithful epitope-focused immunogens.

### Epitope-focused nanoparticle immunization elicits broadly cross-reactive coronavirus antibody responses

We assembled ferritin-based nanoparticles displaying individual epitope-focused scaffold immunogens: ASH1 or ASH2 for the stem helix, and AFP1, AFP3, or AFP11 for the FPPR (***Fig. 2A and Fig. S2A***). These nanoparticles were administered to BALB/c mice as ASH-only, AFP-only, or five-component mixture, with empty nanoparticles as controls (***Fig. 2B and Fig. S2B***). Serum IgG binding was measured against spike trimers from seven human coronaviruses. ASH-based immunization induced preferential reactivity toward SARS-CoV-2 and SARS-CoV spike proteins, with limited binding to seasonal coronaviruses such as NL63 and 229E, consistent with the strong epitope divergence observed across these viruses (***Fig. 1A***). In contrast, AFP-based immunization elicited broader cross-reactivity spanning SARS-CoV-2, SARS-CoV, MERS-CoV, OC43, HKU1 and 229E spike proteins, although responses to NL63 were lower. The mixed immunization regimen produced the broadest overall response profile, combining the strong SARS-related coronavirus reactivity conferred by ASH immunogens with the broader cross-coronavirus recognition elicited by AFP immunogens (***Fig. 2B***).

**Fig. 2.**
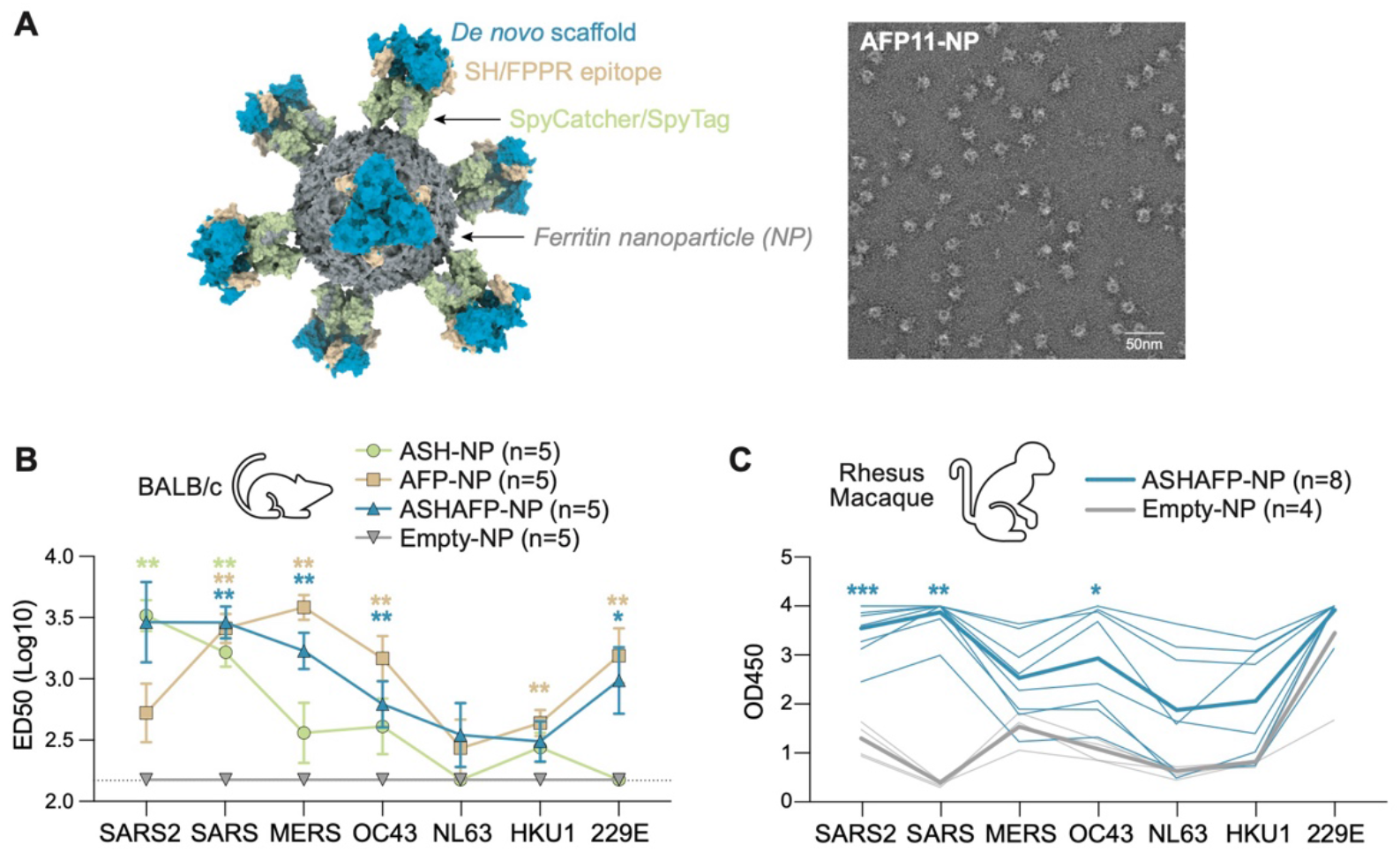
*In vivo* immunogenicity of epitope-focused nanoparticles. **(A)** Schematic and representative negative-stain electron microscopy (NSEM) image of ferritin nanoparticles displaying epitope scaffolds via SpyCatcher/SpyTag conjugation. **(B)** BALB/c mice were immunized with nanoparticle immunogens displaying ASH1 and ASH2 scaffolds (ASH-NP), AFP1, AFP3 and AFP11 scaffolds (AFP-NP), a mixture of ASH-NP and AFP-NP (ASHAFP-NP), or empty nanoparticles (Empty-NP) as a negative control (n = 5 per group). Serum binding to spike proteins from seven human coronaviruses was measured after four doses by ELISA and reported as ED50 values. **(C)** Rhesus macaques were immunized with ASHAFP-NP (n = 8) or empty-NP (n = 4). Serum binding to spike proteins from human coronaviruses was measured after four doses and reported as OD450 values at a 1:200 serum dilution. Data are means ± SEM. Statistical significance was determined by unpaired Mann-Whitney test (*ns*, not significant; \**p* < 0.05; \*\**p* < 0.01; \*\*\**p* < 0.001).

We next evaluated the five-component nanoparticle mixture in rhesus macaques. Immunization induced robust serum binding to SARS-CoV-2 spike, followed by SARS-CoV and OC43 spikes. Antibody responses to MERS-CoV, NL63, and HKU1 spikes were also induced in most animals, with 5 of 8 macaques showing measurable binding, whereas control animals showed minimal reactivity (***Fig. 2C and Fig. S2C***). However, strong baseline reactivity against 229E spike was detected in both vaccine and control groups, precluding assessment of vaccine-associated increases. Such pre-existing serological reactivity may reflect prior exposure to endemic or antigenically related coronaviruses, consistent with previous reports of widespread coronavirus cross-reactive antibody responses in both humans and animal populations (*21, 22*). Collectively, these results demonstrate that co-administration of distinct epitope-focused nanoparticles broadens the landscape of coronavirus cross-reactive antibody responses in both rodent and primate models.

### Epitope-focused vaccination elicits SH- and FPPR-directed broadly neutralizing antibodies

To determine whether the epitope-focused nanoparticles elicited antibodies against the intended S2 epitopes, we isolated antigen-specific B cells from mice immunized with the five-component nanoparticle formulation (***Fig. S3A***). B cells were selected based on dual binding to epitope scaffold probes (ASH and AFP) and spike trimers. Two monoclonal antibodies, 3H7 and 3B2, were recovered and selected for further characterization (***Fig. S3B***). BLI data confirmed that 3H7 and 3B2 targeted the designed SH and FPPR epitopes, respectively. 3H7 bound strongly to SH peptide and SH scaffold (ASH1), with negligible binding to FPPR-derived antigens, whereas 3B2 preferentially recognized FPPR peptide and FPPR scaffold AFP3 (***Fig. 3A***). The antigenic profiles of these vaccine-elicited antibodies closely matched those of the reference bnAbs classes used for immunogen design. The SH-directed antibody 3H7 exhibited a binding pattern characteristic of SH-targeting bnAbs, with strong reactivity toward betacoronaviruses, particularly sarbecoviruses, but minimal binding to alphacoronaviruses 229E and NL63. In contrast, the FPPR-directed antibody 3B2 displayed broader cross-lineage reactivity, including binding to both beta- and alpha-coronaviruses, consistent with the higher conservation of the FPPR epitope (***Fig. 3B and Fig. S3C***).

**Fig. 3.**
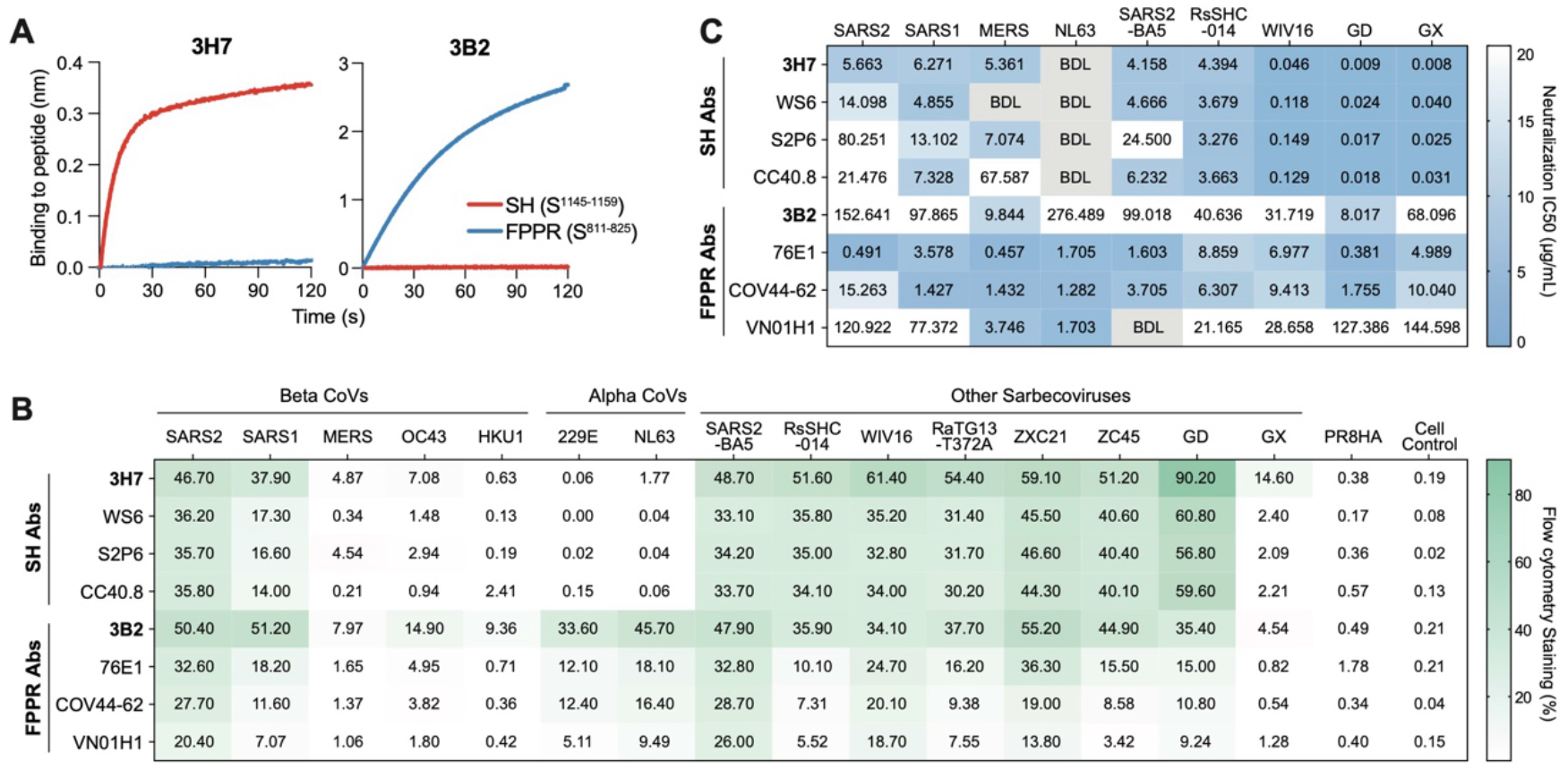
Epitope-focused immunogens precisely elicited antibodies with broad neutralization in mice. MAbs 3H7 and 3B2 were isolated from mice immunized with epitope-focused immunogens. **(A)** Specificity of the isolated mAbs were determined by BLI using SH- and FPPR-derived peptides. **(B)** Neutralizing activity of isolated mAbs against pseudoviruses, reported as half-maximal inhibitory concentration (IC50; μg/mL). BDL, below detection limit, indicates that antibodies failed to achieve 50% neutralization at the highest concentration tested (300 μg/mL). IC50 values were calculated from three independent experiments, each performed in technical duplicate. **(C)** Binding of isolated mAbs to naïve full-length spike proteins expressed on the cell surface, measured by flow cytometry and reported as the percentage of stained live cells.

Neutralization assays further confirmed this correspondence between elicited antibodies and their reference bnAb classes. 3H7 potently neutralized SARS-CoV-2, SARS-CoV, MERS-CoV and multiple sarbecoviruses, demonstrating a robust SH-directed bnAb response. In comparison, 3B2 showed weaker but broader neutralizing activity across selected beta- and alpha-coronaviruses, consistent with the cross-lineage coverage associated with FPPR-directed antibodies (***Fig. 3C and Fig. S3D***). Together, these results demonstrate that epitope-focused vaccination can elicit monoclonal antibodies directed to predefined SH and FPPR bnAb epitopes. Thus, both bnAb epitopes classes independently gave rise to broadly neutralizing antibodies, establishing a closed loop from structurally defined bnAb epitopes, designed scaffold immunogens, to vaccine-elicited antibody responses.

### Heterologous boosting with epitope-focused immunogens protects against distinct coronaviruses

As most individuals have existing SARS-CoV-2 immunity either through vaccination, infection, or both, any broad-based vaccine will need to be tested in a setting of pre-existing SARS-CoV-2 immunity. To model how immunization with our SH and FPPR epitope-focused immunogens boost the widely-prevalent pre-existing immunity, we asked whether epitope-focused nanoparticle boosting could broaden coronavirus immunity in an mRNA-primed setting. BALB/c mice were primed with two doses of SARS-CoV-2 mRNA-LNP vaccine followed by three boosts with ASH-only, AFP-only, or mixed ASH/AFP nanoparticle formulations, matching the designs used in ***Fig. 2A***. Empty nanoparticles served as controls (***Fig. 4A***). Serological analysis showed that mRNA priming dominated antibody responses to SARS-CoV-2 and SARS-CoV spike proteins, with no substantial differences among boosting groups relative to controls. In contrast, AFP-based boosting significantly enhanced antibody binding to multiple heterologous coronavirus spikes, including MERS-CoV, OC43, NL63, HKU1 and 229E. The mixed formulation produced intermediate responses between the ASH and AFP groups, with the most pronounced differences observed for 229E (***Fig. 4B and Fig. S4A***). These results indicate that epitope-focused nanoparticle boosting can reshape antibody breadth beyond the SARS-CoV-2-biased response induced by mRNA priming and expand recognition of divergent coronaviruses.

**Fig. 4.**
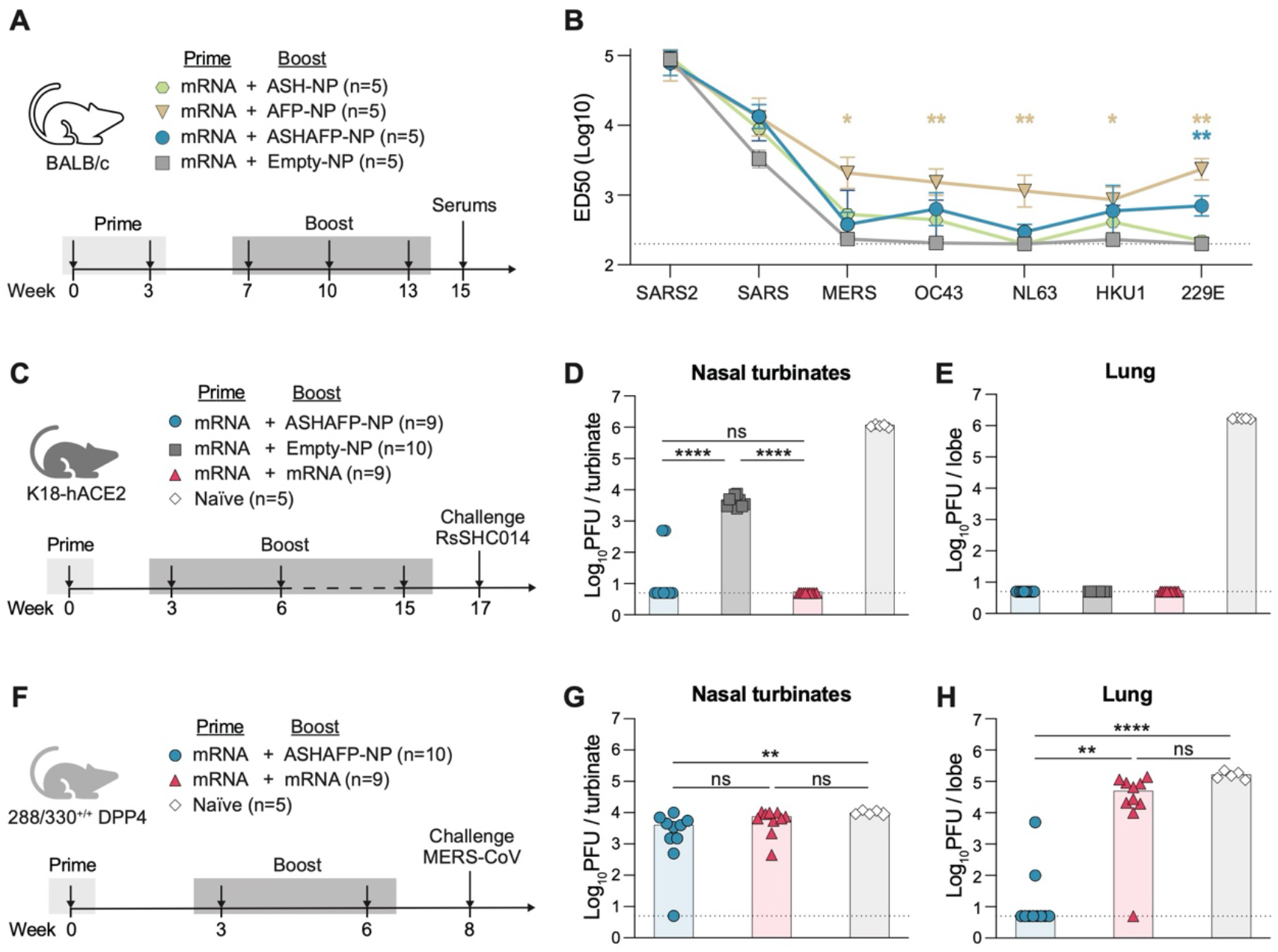
Heterologous boosting protects against authentic SARS-related coronavirus RsSHC014-CoV and MERS-CoV infection in mouse models. Mice were primed with a SARS-CoV-2 mRNA vaccine and boosted with epitope-focused nanoparticle vaccines. **(A) Heterologous prime-boost** regimen in BALB/c mice. **(B)** Serum antibody binding to recombinant spike proteins from human coronaviruses was measured after the final boost by ELISA and reported as ED50 values. **(C)** RsSHC014-CoV challenge study in immunized K18-hACE2 transgenic mice. Viral loads in nasal turbinates **(D)** and lungs **(E)** were quantified by plaque assay of tissue homogenates at 2 days post-inoculation. **(F)** MERS-CoV challenge study in immunized 288/330^+/+^ DPP4 mice. Viral loads in nasal turbinates **(G)** and lungs **(H)** were quantified as above. The plaque assay limit of detection was 5 PFU per tissue. Data are means ± SEM. Statistical significance was determined by unpaired Mann-Whitney test (*ns*, not significant; \**p*< 0.05; \*\**p*< 0.01; \*\*\*\**p*< 0.0001).

To determine whether serological binding translates into protection, we challenged immunized mice with authentic coronaviruses. K18-hACE2 mice were primed with a single dose of SARS-CoV-2 mRNA-LNP vaccine and then boosted three times with ASH/AFP nanoparticles or empty nanoparticles. A parallel group received three additional SARS-CoV-2 mRNA-LNP boosts, and unimmunized mice served as challenge controls. Mice were then challenged with live RsSHC014-CoV, a bat SARS-related zoonotic coronavirus (*23, 24*) (***Fig. 4C***). In the lungs, which reflect lower-airway replication, all SARS-CoV-2 mRNA-LNP–primed groups were strongly protected, with viral titers reduced to undetectable levels in most animals and no significant differences among immunized groups. This protection is consistent with the close antigenic relationship between RsSHC014-CoV and SARS-CoV-2. In the nasal turbinates, which reflect upper-airway replication, both the ASH/AFP nanoparticle-boosted group and the four-dose mRNA-LNP group showed near-complete viral clearance, with viral titers below the limit of detection. By contrast, the mRNA-primed, empty nanoparticle-boosted control group retained high viral loads (***Fig. 4D-E***).

We next assessed cross-protection against MERS-CoV in 288/330^+/+^ DPP4 mice (*25*) using the same prime-boost regimen (***Fig. 4F***). Neutralizing antibodies against MERS-CoV were elicited only by ASH/AFP-nanoparticle boosting, whereas homologous mRNA boosting failed to induce such responses (***Fig. S4B-C***). In the nasal turbinates or the upper respiratory tract, viral loads were comparable across groups (3.80×10^3^, 6.82×10^3^ PFU, 1.01×10^4^ PFU), with only one ASH/AFP-nanoparticle-boosted mouse showing undetectable virus (***Fig. 4G***). In contrast, protection in the lungs or the lower respiratory tract was strongly dependent on the ASH/AFP nanoparticle boost. The three-dose mRNA vaccine group showed no protection and the control group retained high viral titers. By contrast, mice primed with SARS-CoV-2 mRNA-LNP and boosted with ASH/AFP-nanoparticles showed markedly reduced lung viral loads, with most animals near or below the limit of detection (***Fig. 4H***). Together, these challenge studies show that epitope-focused immunogen boosting can harness pre-existing SARS-CoV-2 immunity to confer substantial protection against antigenically distinct coronaviruses, including marked lower-respiratory-tract protection against MERS-CoV.

## DISCUSSION

The rapid antigenic evolution of coronaviruses, combined with the persistent threat of zoonotic spillover from animal reservoirs, makes it impractical to continually update strain-specific vaccines. These challenges underscore the need for immunization strategies that redirect pre-existing immunity, such as widely prevalent infection or vaccine-derived SARS-CoV-2 immunity, toward conserved sites of viral vulnerability. Here, we show that epitope-focused *de novo* design can redirect SARS-CoV-2-specific antibody responses toward the most conserved regions of the coronavirus spike S2 subunit. By presenting the SH and FPPR epitopes on *de novo* designed scaffolds, we elicited antibody responses that closely recapitulate the antigenic and functional features of naturally occurring bnAbs, and conferred broad protection against antigenically divergent coronaviruses.

A major challenge for broad coronavirus vaccine development is that conserved S2 epitopes are typically subdominant during infection or conventional spike vaccination (*4, 6, 11, 12*). In previously vaccinated populations, SARS-CoV-2 mRNA vaccination establishes strong imprinting toward immunodominant Spike epitopes, which may limit responses to more conserved regions (*26*). Broad vaccine strategies must therefore either overcome this imprinting or exploit it by redirecting pre-existing immunity. In this setting, boosting with SH and FPPR epitope-focused immunogens reshaped pre-existing SARS-CoV-2 mRNA-LNP vaccine-induced responses, selectively enhancing antibody recognition of antigenically distant coronaviruses while preserving responses to closely related sarbecoviruses. This effect was particularly evident in the enhanced serological breadth and *in vivo* protection against MERS-CoV, a phylogenetically distant betacoronavirus that is not protected by SARS-CoV-2-based immunity alone (*27*). These findings indicate that conserved S2 epitopes can be leveraged to extend protection across major evolutionary branches of the betacoronavirus family.

The concept of focusing antibody responses on conserved but subdominant neutralizing epitopes has been explored for several rapidly evolving viruses (*28*), including influenza viruses (*29-31*), flaviviruses (*32*), respiratory syncytial virus (RSV) (*33-35*), and HIV-1 (*36-39*). Substantial efforts have been devoted to focusing immune responses toward conserved neutralizing sites through antigen engineering (*40*), epitope masking (*41*), germline targeting (*42*), or stabilized immunogen design (*29, 33*). Epitope-focused vaccine strategies have further evolved through structure-guided epitope grafting and stabilized scaffold-based approaches to recapitulate the antigenic features of bnAb epitopes (*28, 34*). Early studies in HIV-1 demonstrated that structurally defined epitopes could be transplanted onto heterolo-gous scaffolds to engage desired antibody lineages, providing a conceptual basis for epitope-centric vaccine design (*28*). Similar principles have been extended to RSV and influenza virus, where stabilization of conserved fusion machinery, including prefusion RSV F protein (*34*) and influenza hemagglutinin stem (*29*), has enabled partial redirection of immune responses toward bnAb determinants.

For coronaviruses, previous efforts to target conserved S2 epitopes have included peptide immunization (*43*), engineered S2 subunit antigens (*21, 44*), and scaffold-based grafting of stem helix or fusion-associated regions (*15*). Although these approaches have provided evidence that conserved S2 elements are antigenically accessible, they have generally shown limited success in reproducibly eliciting bnAb responses and have not demonstrated protection against *in vivo* challenge. Linear peptides may fail to maintain the conformational constraints required to faithfully mimic native epitope geometry (*45*), whereas larger S2-derived antigens may retain structural flexibility or competing immunodominant surfaces that reduce epitope focusing. In contrast, the *de novo* design strategy used here enabled precise structural stabilization of isolated SH and FPPR epitopes in conformations that closely match their anti-body-bound states, while minimizing unrelated antigenic features. The resulting immunogens exhibited strong antigenic agreement with reference bnAbs and reproducibly elicited antibody responses with matched specificity and function, supporting the importance of native-like structural presentation for induction of bnAb-like immunity.

These findings suggest that scaffold-based epitope focusing may provide a generalizable strategy for vaccine design against antigenically variable pathogens. By extracting conserved neutralizing epitopes from complex viral surface proteins and presenting them in optimized structural contexts, this approach may selectively amplify broadly protective antibody responses that are otherwise inefficiently induced by infection or conventional vaccination. The concordance observed here, where reference bnAbs defined the target epitope conformations, designed scaffolds faithfully presented those epitopes, and vaccination elicited antibodies with matching antigenic profiles, further supports the feasibility of extending this strategy to other pathogens for which conserved vulnerable epitopes have already been structurally and antigenically defined.

Together, our findings demonstrate that *de novo* epitope-focused immunogen design can reproducibly elicit protective immunity against conserved but immunologically subdominant coronavirus epitopes and can harness prior SARS-CoV-2 immunity towards protective immunity. By redirecting antibody responses toward multiple conserved S2 epitopes, this strategy broadens the range of targetable sites for pan-coronavirus vaccine design beyond conven-tional spike-based immunization approaches. The protective efficacy of these immunogens as heterologous boosts also provides a practical translational path for enhancing existing vaccine platforms in populations with prior SARS-CoV-2 immunity. More broadly, this work supports the integration of structural vaccinology with AI-enabled protein design as a general strategy for engineering immune specificity at the epitope level, with potential applications in the development of next-generation broad-spectrum vaccines against rapidly evolving pathogens.

## MATERIALS AND METHODS

Detailed materials and methods are provided in Supplementary Materials.

## ACKNOWLEDGMENTS

We thank Walvax Biotechnology Co., Ltd. for providing the mRNA vaccine; Dr. Zhiqiang Ku (Westlake University) for providing the expression plasmids of antibody 76E1, COV44-62 and VN01H1; Dr. Qiang Sun and Dr. Yong Lu (Institute of Neuroscience, Chinese Academy of Sciences) for the help with the monkey experiments; the technical support from the Biomedical Research Core Facility, Laboratory Animal Resources Center, and High-Performance Computing Center at Westlake University.

## Funding

Westlake Education Foundation (D.L.)

Westlake University Research Center for Industries of the Future (D.L.)

Zhejiang Provincial Key Laboratory Construction Project 2024ZY01026 (D.L.)

The National Natural Science Foundation of China 82471858 (D.L.)

Zhejiang Provincial Natural Science Foundation of China LR26C080001 (D.L.)

China Postdoctoral Science Foundation 2023M733182 (M.L.)

The National Natural Science Foundation of China 82402131 (M.L.)

Major Project of Guangzhou National Laboratory GZNL2024A01017 (X.M.)

Hanna H. Gray Fellowship from the Howard Hughes Medical institute (D.M.)

## Author contributions

Conceptualization: D.L. and D.R.M. Methodology: M.L., J.S., F.Z.P., Z.Z., Z.W.

Investigation: M.L., Q.C., J.S., Y.L., F.Z.P, K.H., Z.D., C.C.

Visualization: D.L., M.L., Z.D., D.R.M.

Funding acquisition: D.L., D.R.M., X.M., M.L.

Project administration: D.L., D.R.M., X.M., M.L.

Supervision: D.L., D.R.M., X.M., F.Y., M.L.

Writing – original draft: D.L. and M.L.

Writing – review & editing: D.L., D.R.M., X.M.

## Data, code, and materials availability

All data associated with this manuscript including the epitope scaffolds and antibody sequences are available in the main text or the supplementary materials (***Table S1***). Materials described in this manuscript will be available through a material transfer agreement with Westlake University.

## Supplementary Materials include

Materials and Methods

Figs. S1 to S4

Table S1

