## Supplementary Materials for "Epitope-Focused Immunogens Confer Broad Protection against Coronaviruses"

Mingxi Li, et al.

**This file includes:**

Materials and Methods

Supplementary Figures and Figure Legends (Fig. S1-S4)

Supplementary Table

### MATERIALS AND METHODS

#### Computational design of epitope scaffold immunogens

**De novo design of epitope scaffolds.** The stem helix (SH) epitope $^{1146}$DSFKEELDKYF KNH$^{1159}$ from PDB: 7RNJ and fusion peptide proximal region (FPPR) epitope $^{813}$SKRFIEDL LFNK$^{825}$ from PDB: 7X9E were extracted from experimentally determined coronavirus spike structures and used as independent structural motifs for de novo scaffold design. SH- and FPPR-based immunogens were designed separately using the RFdiffusion protein design framework in scaffold generation mode. During the diffusion process, the target epitope was treated as a fixed structural motif, with both backbone and side-chain conformations constrained to preserve the native epitope geometry observed in the parental spike protein. To maximize epitope accessibility, design constraints were imposed such that the target epitope remained solvent exposed on the surface of the scaffold, while the surrounding scaffold architecture avoided steric occlusion of the antibody-accessible surface.

Scaffold proteins were designed with lengths ranging from 100 to 150 amino acids to balance structural stability, independent folding, recombinant expression, and sufficient structural support for the constrained epitope. Approximately 10,000 scaffold backbones were generated independently for each target epitope.

For each RFdiffusion-generated backbone, amino acid sequences were designed using ProteinMPNN. Eight candidate sequences were generated for each backbone, and the highest-scoring sequence according to the ProteinMPNN model was selected for subsequent computational evaluation.

**In silico screening of designed scaffold proteins.** Designed scaffold proteins were subjected to a multi-step computational screening pipeline to evaluate structural stability, epitope preservation, and compatibility with broadly neutralizing antibody recognition.

First, all designed sequences were structurally re-predicted using AlphaFold3. Candidates with an average predicted local distance difference test (pLDDT) score greater than 80 were retained as high-confidence folded structures. The AlphaFold3-predicted structures were subsequently aligned to their corresponding design models, and backbone root-mean-square deviation (RMSD) was calculated over the constrained epitope residues. Designs exhibiting epitope backbone RMSD values below 2 Å were selected for further analysis, indicating successful preservation of the intended epitope geometry following sequence optimization.

To evaluate antibody accessibility, the remaining scaffold candidates were analyzed at the complex level by structural prediction with their corresponding broadly neutralizing antibodies. Predicted antibody–scaffold complexes were assessed using interface quality metrics, including interface geometry and steric clash scores. Designs exhibiting substantial steric interference, poor interface complementarity, or scaffold-mediated obstruction of antibody access to the target epitope were excluded. This screening strategy enriched for scaffold proteins that not only maintained the native conformation of the target epitope but also accurately recapitulated its antibody-recognizable structural presentation.

**Recombinant protein expression and purification**

**Epitope scaffolds.** Codon-optimized sequences encoding the epitope scaffolds were commercially synthesized and individually cloned into pVRC8400 (SinbioB) with a C-terminal 6×His-tag. Recombinant proteins were transiently expressed in Expi293F cells with PEI MAX (Polysciences). Supernatants were collected 5 days post-transfection, clarified by centrifugation, filtered through a 0.22-μm membrane, and purified by $Ni^{2+}$ affinity chromatography (SunResin, Cat # A453201) using an ÄKTA system and further purified by size exclusion chromatography on Superdex 200 Increase 10/300 GL.

**Monoclonal antibodies.** Heavy- and light-chain plasmids were co-transfected into Expi293F cells at a 1:1.5 ratio using PEI MAX. Supernatants were collected 5 days post-transfection, clarified by centrifugation, filtered through a 0.22-μm membrane. Human monoclonal antibodies were purified by protein A affinity chromatography (SunResin, Cat # A4093101) and mouse monoclonal antibodies were purified by protein G affinity chromatography (Genscript, Cat # L00681) on an ÄKTA system. Bound antibodies were eluted with 100 mM glycine (pH 2.5), immediately neutralized with 1/10 volume of 1 M Tris-HCl, and buffer-exchanged into PBS.

**Nanoparticles.** *Helicobacter pylori* ferritin with N96 and SpyTag at N-terminus and 6×His-tag at C-terminus(*1*) was expressed in Expi293F cells with PEI MAX. Supernatants were collected 5 days post-transfection, clarified by centrifugation, filtered through a 0.22-μm membrane, and purified by HisTrap HP affinity chromatography (Cytiva, Cat # 17524801) followed by size exclusion chromatography on Superose 6 Increase. Epitope scaffold-SpyCatchers were expressed as a fusion protein with a C-terminal 6×His-tag and purified using $Ni^{2+}$ affinity chromatography followed by size exclusion chromatography on Superdex 200 Increase 10/300 GL. A 1:1 molar ratio of ferritin-SpyTag and immunogen-SpyCatcher components were combined and incubated at 4**°C** overnight followed by size exclusion column on Superose 6 Increase in PBS to separate conjugated products from residual components. The conjugated nanoparticle product was then run through SDS-PAGE to verify conjugation and analyzed by negative-stain EM.

**Spike trimers.** To generate soluble S ectodomain proteins from SARS-CoV-2 (residues 1-1208; GenBank: MN908947.3), SARS-CoV-1 (residues 1-1190; GenBank: ABD72970.1), MERS-CoV (residues 1-1291; GenBank: AHI48572.1), HCoV-HKU1 (residue 1-1295; GenBank: URC25188.1), HCoV-OC43 (residues 1-1300; GenBank: AVR40344.1), HCoV-229E (residues 1-1110; GenBank: APT69883.1) and HCoV-NL63 (residues 1-1291; GenBank: ARU07594.1), we synthesized the DNA fragments from Genscript and cloned them into the pCAGGS vector. Double proline substitutions (2P) were introduced into the S2 subunit: K968/V969 in SARS-CoV-1, K986/V987 in SARS-CoV-2, V1060/L1061 in MERS-CoV, A1071/L1072 in HCoV-HKU1, A1078/L1079 in HCoV-OC43, S1052/I1053 in HCoV-NL63 and T871/I872 in HCoV-229E were replaced by proline. The furin cleavage sites of SARS-CoV-2 were replaced by a "GSAS" linker. The trimerization T4 fibritin motif was incorporated at the C-terminus of the S proteins. To purify and biotinylate the spike proteins, 8x HisTag and AviTag spaced by GS-linkers were added to the C-terminus after the trimerization motif. Recombinant proteins were transiently expressed in Expi293F cells with PEI MAX (Polysciences). Supernatants were collected 5 days post-transfection, clarified by centrifugation, filtered through a 0.22-μm membrane, and purified by HisTrap HP affinity chromatography (Cytiva, Cat # 17524801) followed by size exclusion chromatography on Superdex 200 Increase 10/300 GL.

**ISCOM preparation**

Solutions of cholesterol (20mg/ml; sigma, 700000P) and DPPC (1,2-dipalmitoyl-sn-glycero-3-phsophocholine (20 mg/ml, sigma, 850355P) were prepared in Milli-Q water contain 20% (w/v) MEGA-10 (Sigma, 850542P) detergent at 60°C. Quil-A saponin was dissolved in Milli-Q water at a final concentration of 100 mg/ml. All components were mixed at 60°C at a molar ratio of 10:10:5 (Quil-A:cholesterol:DPPC) followed by dilution with PBS to a final concentration of cholesterol at 1 mg/ml. The solution was allowed to equilibrate overnight at room temperature, followed by dialysis against PBS using a 12,000 MWO (molecular weight cut-off, sigma, PURX12050) membrane for 5days (change PBS twice a day). The adjuvant solution was then concentrated 10-fold using a 50,000 MWCO ultrafiltration tube (Millipore, UFC805024). Final concentration of ISCOM was recorded as the Quil-A concentration in the solution determined by the sulfuric acid-phenol method [29].

**Surface plasmon resonance**

Binding kinetics between epitope scaffolds and target antibodies was measured by Surface Plasmon Resonance (SPR) on a Biacore 8K+ instrument using Protein-A coated S series chips. The target antibodies were individually captured for 60 sec at 10 μL/min to a level of 1000-2000RU onto this surface. The epitope scaffolds were subsequently injected as analytes at five or eight concentrations using the single cycle injection method. The association phase was carried out for 120 seconds and the dissociation was done for 300 seconds with PBST buffer flowing at 30 μL/min. Regeneration of the binding surface was done in 10 mM glycine-HC, pH=2 for 30 seconds at 30 μL/ min with a 30 second baseline stabilization. A 1:1 Langmuir or Heterogenous ligand model was used for data fitting and analysis.

**Sera binding by ELISA**

96-well high-binding ELISA plates were coated overnight at 4 °C with SARS-CoV-2 spike protein (1 μg/mL). Plates were washed three times with PBST (0.1% Tween-20 in PBS) and blocked with 3% BSA in PBST for 2 h at 37 °C. Human sera were diluted at 1:100 and incubated for 1 h at 37 °C. Mouse sera were serially diluted 3-fold from an initial dilution ratio of 1:200 and incubated for 1 h at 37 °C. Rhesus macaque sera were diluted at 1:200 and incubated for 1 h at 37 °C. After five washes, plates were incubated with HRP-conjugated anti-human IgG antibody (Promega, Cat # W4031; 1:5,000 dilution) or HRP-conjugated anti-mouse IgG antibody (Promega, Cat # W4021; 1:5,000 dilution) for 1 h at 37 °C. Plates were washed five times, developed with TMB substrate (Cwbio, Cat # CW0050) for 15 min at room temperature in the dark, and the reaction was terminated with 50 μL of 1 M sulfuric acid. Absorbance was measured at 450 nm.

**Mouse immunization and protection studies**

**Homologous immunization of nanoparticles.** Female BALB/c mice aged 6–8 weeks were purchased from Charles River and randomly divided into 4 groups (n = 5 in each group). Mice were vaccinated intramuscularly with ASH1 and ASH2-NP mixture; AFP1, AFP3 and AFP11-NP mixture; five-component mixture; empty NP. Nanoparticles were administered at 10 μg in 50 μL PBS and formulated with 50 μL AddaVAX (InvivoGen, Cat # vac-adx-10). Mice were immunized at week 0, 3, 6 and 9. Sera were collected 2 weeks after $4^{th}$ immunization, heat-inactivated at 56°C for 30min, and stored at -80°C before analysis.

**Heterologous immunization of mRNA and nanoparticles.** Female BALB/c mice aged 6–8 weeks were purchased from Charles River and randomly divided into 4 groups (n = 5 in each group). Mice were vaccinated intramuscularly twice with mRNA RQ3033 (Omicron XBB.1.5 spike) from Walvax Biotechnology Co., Ltd. followed by three additional doses of nanoparticles (10 μg in 50 μL PBS and 50 μL AddaVAX). Mice were immunized at week 0, 3, 7, 10 and 13. Sera were collected 2 weeks after the last immunization, heat-inactivated at 56°C for 30min, and stored at -80°C before analysis.

**Mouse challenge studies.** K18-hACE2 mice (B6. Cg-Tg (K18-ACE2) 2 Prlmn/J; JAX strain number #034860) were used for the RsSHC014-CoV challenge study, and hDPP4-KI mice with an altered DPP4 at amino acid positions 288 and 330 were bred and used for the MERS-CoV challenge study (*2*). Mice were vaccinated intramuscularly once with 1 μg of SARS-CoV-2 mRNA-LNP (Comirnaty (BNT162b2) mRNA-LNP, Pfizer-BioNTech), followed by three or two additional doses of either nanoparticles or mRNA-LNP. A naïve group was included for comparison. All mice were anesthetized by using 30% isoflurane diluted in propylene glycol and infected intranasally with RsSHC014 ($1 \times 10^5$ PFU/ml) or MERS-CoV ($1 \times 10^5$ PFU/ml), which have been described previously (*2*). Viral loads in nasal turbinates and lungs from each vaccination group were quantified by plaque assay of tissue homogenates at 2 days post-infection (dpi) which is peak replication.

**Rhesus macaque immunization.** Twelve adult rhesus macaques aged between 5 and 9 years were intramuscularly vaccinated with 50 μg five-component NP mixture in 2mL PBS and formulated with 50 μg ISCOM adjuvant. Macaques were immunized at week 0, 4, 8 and 12. Sera were collected 2 weeks after $4^{th}$ immunization, heat-inactivated at 56°C for 30min, and stored at -80°C before analysis.

**Pseudovirus neutralization assay**

Neutralizing activity of antibodies were determined using pseudovirus as previously reported (*3*). Briefly, antibodies were 3-fold serially diluted in 96-well cell culture plates, mixed with pseudovirus, and incubated at 37°C for 1 h. Huh7 cells for MERS and 293T-ACE2-TMPRSS2 cells for other coronaviruses were then added to the mixture of antibody-pseudovirus, incubated at 37°C for additional 48 h, and lysed for measuring luciferase-activity. The IC50 values were calculated based on the reduction of 50% relative light units (Bright-Lite Luciferase Assay System, Vazyme, Cat # DD1204) compared to the virus-only control, by fitting dose-response curves with an asymmetric five-parameter nonlinear regression model in GraphPad Prism 10.

**Isolation of antigen-specific single B cells by FACS**

Antigen-specific single B cells were sorted as previously described(*4*). In brief, single splenocyte suspension from mice vaccinated with five-component nanoparticle mixture; were collected and incubated with an antibody and antigen cocktail for identification of specific B cells. The antibody cocktail consisted of CD3- PerCP/Cy5.5, B220-APC/Cy7, CD19-AF700, IgG1-FITC, IgG2a-FITC, IgG2b-FITC, IgG3-FITC. The epitope-scaffold probe pool was labeled by Streptavidin-APC, containing ASH1, ASH2, AFP1, AFP3 and AFP11. The spike probe pool was labeled by Streptavidin-PE, containing SARS-CoV-2, SARS-CoV, MERS-CoV and OC43 spike proteins. The scaffold-only probe pool was labeled by Streptavidin-BV421, containing the empty scaffolds from ASH1, AFP1, AFP3 and AFP11. Dead cells were stained using the Zombie Aqua Fixable Viability Kit (BioLegend, Cat # 423102). The dual-binding single B cells were gated as B220+ CD19+ IgG+ Scaffold- Epitope scaffold+ Spike+ and sorted into 96-well PCR plates containing 19ul 0.2% triton-100(sigma) and 1ul Recombinant RNase Inhibitor (TaKaRa, Cat # 2313A). Plates were then snap-frozen on dry ice and stored at -80°C until RT reaction. All cytometric data were collected on CytoFLEX SRT cytometer (Beckman). The gating strategies of flow cytometry are shown in Figure S3A.

**Single B cell PCR, cloning, and expression of mAbs**

Single B-cell antibody gene amplification and cloning were performed as previously described(*4*). Briefly, reverse transcription and cDNA amplification was performed using standard smart-seq2 protocol using the HiScript III First-Strand cDNA Synthesis Kit (Vazyme, Cat # R312). Then the purified cDNA of each cell was used for the following BCR target enrichment by a 2-step nest PCR strategy using IS primer and primers targeting the 5′ leader and immunoglobulin constant regions. The IgG heavy and light chain variable genes were amplified by nested PCR and cloned into expression vectors to produce full mouse IgG1 antibodies.

**Bio-Layer Interferometry (BLI)**

Binding ability were characterized at 30 °C using an Octet R8 system (Sartorius). Peptide SH (LDSFKEELDKYFKNH) and FPPR (KPSKRSFIEDLLFNK) were synthesized with biotin at N-terminus (QYAOBIO). For epitope screening, peptide was immobilized onto Streptavidin biosensors (Sartorius, Cat# 18-5019) in assay buffer (PBS + 0.1% BSA) to a level of 0.2 nm. Then, the sensors were exposed to mAb for 120s to monitor binding. Data were processed in Octet Analysis Studio.

**Cell surface staining**

HEK 293T cells were transfected with expression plasmids encoding coronavirus spikes, and incubated at 37 °C for 24 h. Cells were digested from the plate with TrypLE Express (Thermo, Cat # 12604021) and distributed onto 96-well plates. Cells were washed twice with 200 µL staining buffer (PBS with 1% heated-inactivated fetal bovine serum (FBS)) between each of the following steps. First, cells were stained with each monomeric antibody (10 µg/mL) at 4 °C for 30 min. Anti-Human IgG-APC (Biolegend, Cat# 410712, 1:200 dilution) or anti-mouse IgG1-APC (Biolegend, Cat# 406610, 1:200 dilution) were added and incubated at 4 °C for 20 min. After extensive washes, the cells were resuspended and analyzed by Cytek Aurora and FlowJo 10 software. HEK 293T cells with mock transfection were stained as background control.

**Production of pseudoviruses**

Pseudoviruses carrying the full-length spike envelope of coronaviruses were generated as previously reported (*3*). Specifically, human immunodeficiency virus backbones expressing firefly luciferase (pNL4-3-R-E-luciferase) and pcDNA3.1 vector encoding either SARS-CoV-2 or coronavirus spike proteins were co-transfected into the HEK-293T cells (ATCC). Forty-eight hours later, pseudoviruses in the viral supernatant were collected, centrifuged to remove cell lysis, and stored at -80°C until use. The SARS2 was the prototype strain (GenBank: MN908947.3) with a D614G mutation and the Omicron BA.4/5 variant was constructed (Pango lineage BA.4/5, GISAID: EPI_ISL_12559461). The cDNAs encoding the SARS-CoV-1 spike (NCBI Accession NP_828851.1), MERS-CoV spike (GenBank: AHI48572.1), NL63-CoV spike (GenBank: ARU07594.1), Bat SARS-like coronavirus

RsSHC014 (GenBank: AGZ48806.1), Bat SARS-like coronavirus WIV16 (GenBank: ALK02457.1), Pangolin CoV GD spike (GenBank: QLR06867.1) and Pangolin CoV GX spike (GenBank: QIA48614.1) were synthesized with codons optimized for protein expression (Genscript) and verified by sequencing. NL63-CoV spike was mutated with A685V, L853F and Q998K (*5*) and RsSHC014 spike was mutated with Y623H (*6*) for more infection of pseudoviruses.

**Circular dichroism**

Far-ultraviolet CD measurements were carried out with Chirascan V100 (Applied Photophysics) equipped with a temperature-controlled multi-cell holder. Wavelength scans were measured from 260 to 190 nm at 25, 95°C and again at 25°C after fast refolding. Temperature melts monitored dichroism signal at 222 nm in steps of 2°C/minute with 30s of equilibration time. Wavelength scans and temperature melts were performed using 0.3 mg/ml protein in PBS buffer with a 1 mm path-length cuvette. Melting temperatures were determined fitting the data with a sigmoid curve equation. All designs retained more than half of the mean residue ellipticity values, which indicated the Tm values are greater than 95°C.

**Negative-stain electron microscopy (NSEM)**

Samples were diluted to 0.02–0.05 mg/ml with a buffer containing 10mM Tris-HCl, pH 8, and 500 mM NaCl. A 2.5-µl drop of the diluted sample was applied to a glow-discharged carbon-coated copper grid for approximately 60 s. The drop was removed using filter paper, and the grid was washed with a 2.5-µl drop of 3% uranyl acetate, then removed it with filter paper immediately. Adsorbed proteins were negatively stained by applying a 2.5-µl drop of 3% uranyl acetate for 30s and removing it with filter paper. Micrographs were collected using Talos L120C microscope (ThermoFisher) operating at an acceleration voltage of 120kV and equipped with a Ceta CCD camera. Images were recorded at a magnification of 57000x and a defocus of 2.5 µm.

**Ethics statement**

Mouse immunization and characterization were conducted in the animal facility of Westlake University and Yale University. All experiments conducted in the animal facility of Westlake University were carried out in strict compliance with the Guide for the Care and Use of Laboratory Animals of the People's Republic of China and approved by the Committee on the Ethics of Animal Experiments of Westlake University (AP#22-031-3-LDP-3).

Mouse infections were performed in an animal biosafety level 3 (ABSL-3) facility with approval from the Yale Institutional Animal Care and Use Committee (IACUC 2024-20520) and Yale Environmental Health and Safety.

Rhesus macaque experiments were performed in the animal facility of Institute of Neuroscience, Chinses Academy of Sciences, China, and approved by the Primate Life Sciences Ethics Committee of the Centre for Excellence in Brain Science and Intelligence Technology, Chinese Academy of Sciences (CEBSIT-2023047). In line with reduction principles, vaccination and blood collection procedures were conducted with utmost care to minimize pain and discomfort. Convalescent blood samples were collected from 38 donors previously infected or vaccinated with SARS-CoV-2. Serum samples were obtained. This study was approved by the Ethics Review Committees of Westlake University (20220410LDP001).

**Quantification and Statistical Analysis**

GraphPad Prism 10 software was used for data visualization and statistical analyses. Statistical comparisons between two groups were performed using unpaired, two-tailed Mann-Whitney test. Data are presented as mean ± SEM unless otherwise indicated. Statistical significance was defined as ns (not significant, $p \geq 0.05$), * $p < 0.05$, ** $p < 0.01$ and *** $p < 0.001$.

### SUPPLEMENTARY FIGURES AND FIGURE LEGENDS

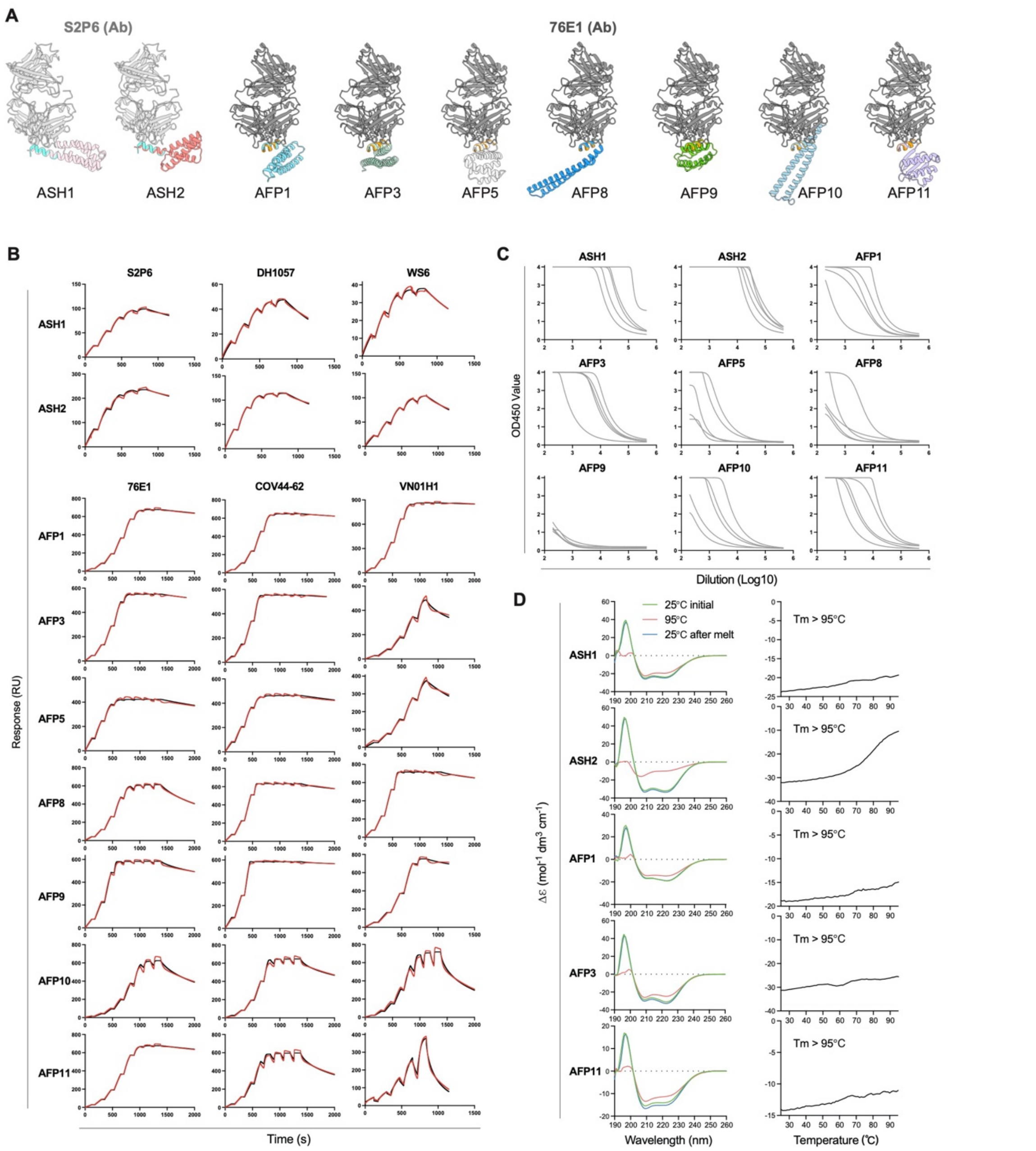


**Fig. S1. Characterization of epitope scaffolds. (A)** Conformations of the SH epitope (cyan) and FPPR epitope (orange) presented on epitope scaffolds and bound by S2P6 and 76E1 mAbs, respectively. Reference antibody–epitope structures are from PDB 7RNJ and 7X9E. **(B)** SPR binding curves of representative SH- and FPPR-directed mAbs to epitope scaffolds. Acquired data are shown in black, and fitted curves are shown in red. Binding affinities are reported as *K*D values in Fig. 1D. Data are representative of at least two independent experiments. **(C)** ELISA binding of mouse sera to SARS-CoV-2 Spike. Sera were collected from BALB/c mice immunized with epitope-scaffold monomers. Data are means from two independent experiments; corresponding ED50 values are summarized in Fig. 1E. **(D)** Circular dichroism (CD) spectra of epitope scaffolds measured at different temperatures, with CD signal at 222 nm plotted as a function of temperature.

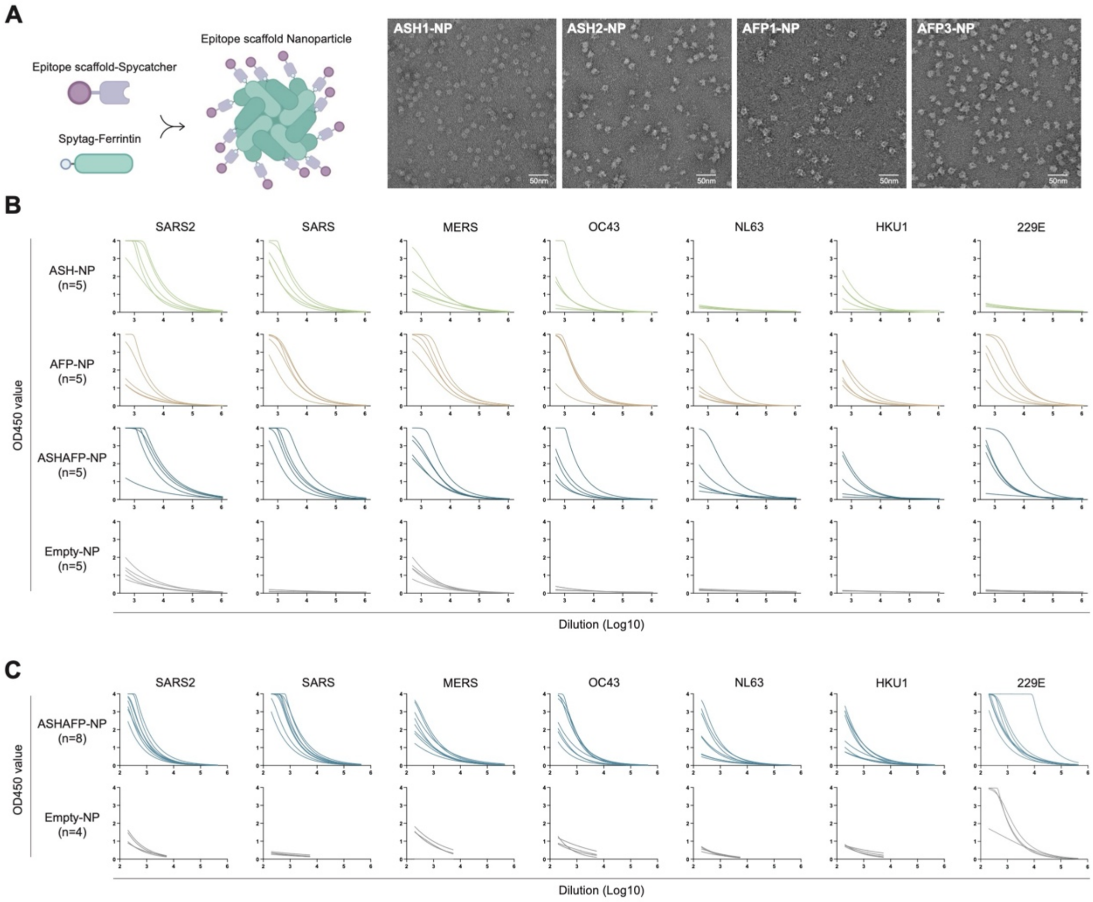


**Fig. S2. Serum binding after immunization with epitope-focused nanoparticles. (A)** Design and negative-stain electron microscopy (NSEM) analysis of SpyCatcher/SpyTag ferritin nanoparticles displaying 24 copies of epitope scaffolds. **(B,C)** Representative ELISA binding curves of sera from immunized BALB/c mice **(B)** or rhesus macaques **(C)** against Spike proteins from human coronaviruses. Data are means from two independent experiments; corresponding ED50 values or OD450 values at a 1:200 dilution are summarized in Fig. 2.

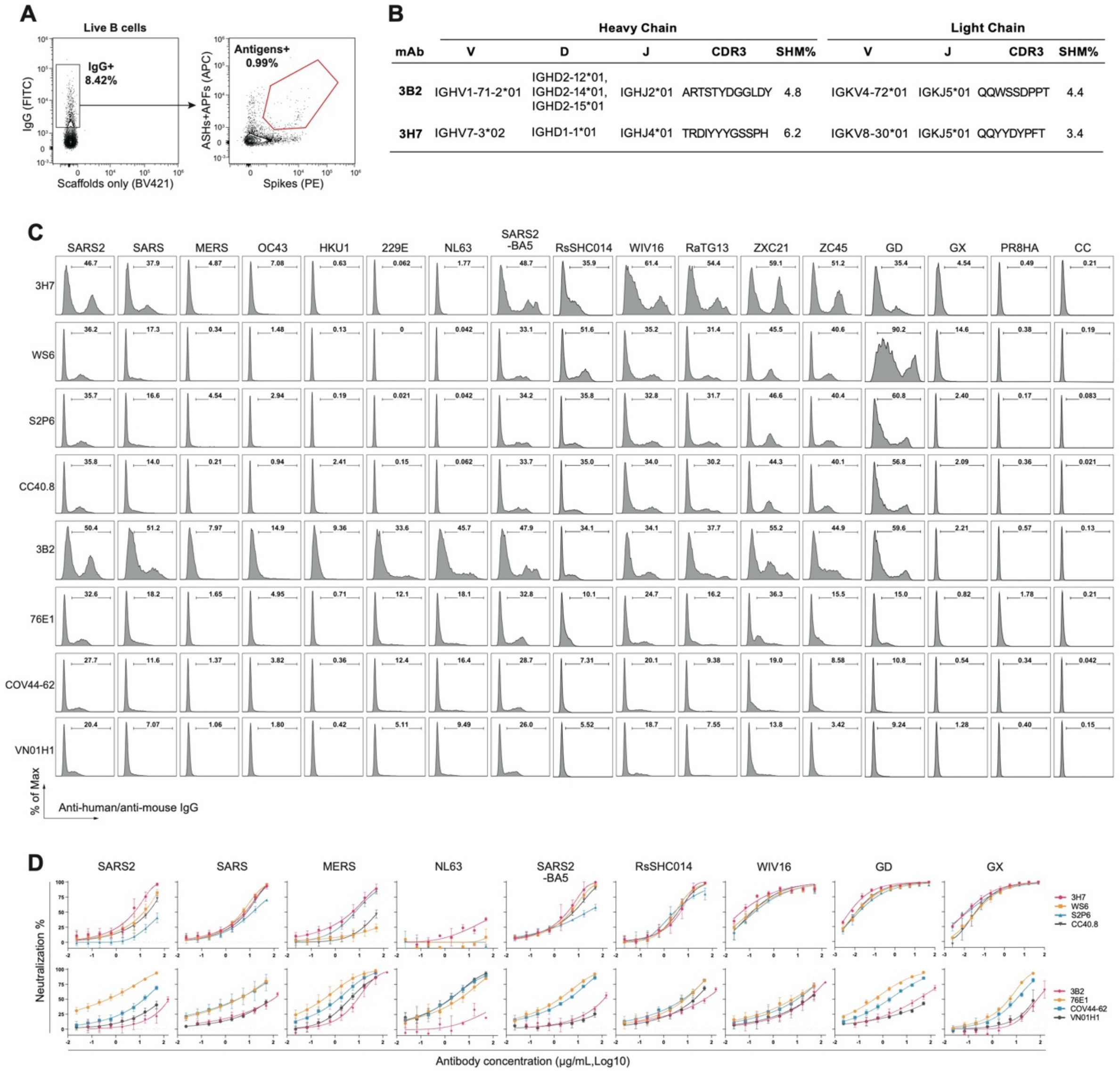


**Fig. S3. Isolation of antigen-specific antibodies from epitope-focused nanoparticles-immunized mice. (A)** Representative flow-cytometric gating strategy for single-cell sorting of antigen-specific IgG+ B cells recognizing both epitope scaffolds and Spike proteins. The epitope-scaffold probe pool contained ASH1, ASH2, AFP1, AFP3 and AFP11. The Spike probe pool contained SARS-CoV-2, SARS-CoV, MERS-CoV and OC43 Spike proteins. **(B)** Immunogenetic features of two neutralizing BCR sequences, including V/J gene usage and alleles, heavy- and light-chain CDR3 amino acid sequences, and somatic hypermutation frequencies. **(C)** Representative flow-cytometric analysis of antibody binding to full-length Spike proteins. HEK293T cells expressing the indicated Spike proteins were incubated with individual antibodies, followed by staining with fluorophore-conjugated secondary antibodies. Influenza H1N1 PR8 hemagglutinin (PR8HA) and untransfected cells (CC) were used as negative controls. Related to Fig. 3B. **(D)** Representative neutralization curves of isolated mAbs and control mAbs against pseudoviruses. Data are means ± SEM from three independent experiments; corresponding IC50 values are summarized in Fig. 3C.

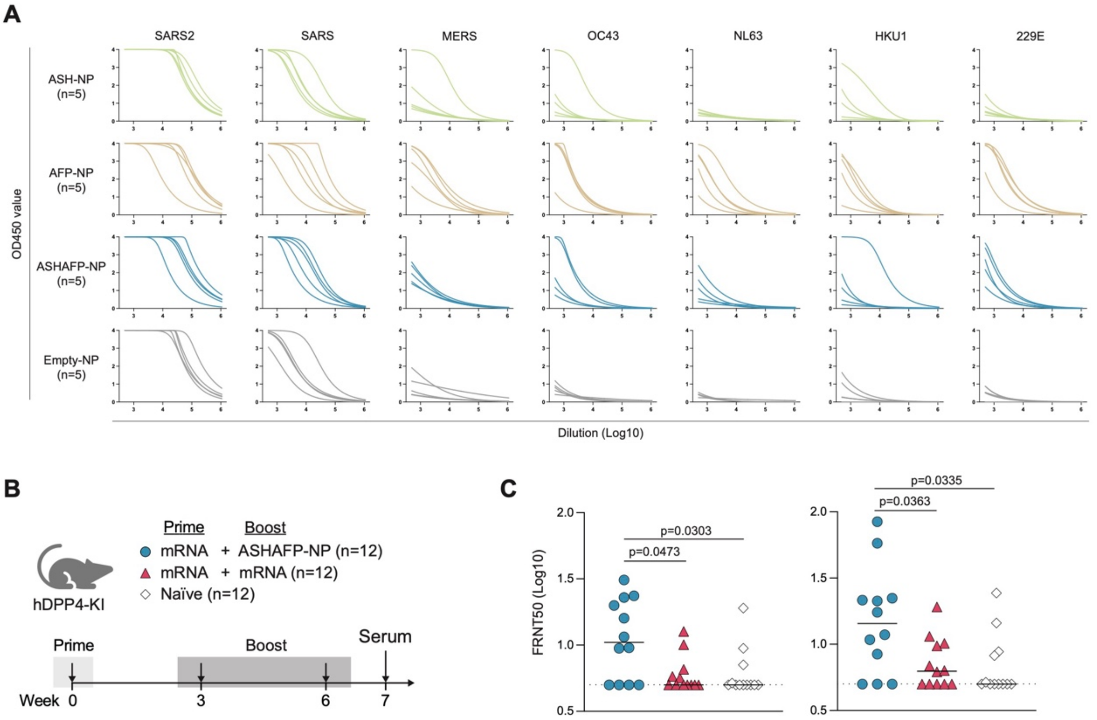


**Fig. S4. Serum binding and neutralization after heterologous boosting. (A)** Representative ELISA binding curves of sera from BALB/c mice immunized with SARS-CoV-2 mRNA prime followed by nanoparticle boosts. Data are means from two independent experiments; corresponding ED50 values are summarized in Fig. 4B. **(B)** Vaccination and serum-collection schedule for hDPP4-KI mice at 7 weeks post-prime. **(C)** Serum neutralization of authentic MERS-CoV was measured by plaque reduction neutralization test and reported as FRNT50 (50% focus reduction neutralization titer) values. Data from two independent experiments are shown. Statistical significance was determined by unpaired Mann-Whitney test.

### SUPPLEMENTARY TABLE

| Epitope scaffold | Amino acid sequence |
|---|---|
| ASH1 | DSFKEELDKYFKNHAAVKATAEAAIAAFDAAAAAGDYAAAAAALAQLVALFNKYKDDPANRAI LLTLYATLLAVRAAHPEAAADPAVAAEWAELQAAAAA |
| ASH2 | DSFKEELDKYFKNHIWIIIAEELAEKCKKAAENPDPEFVKKVREYFKEVKELVEKLKPKLPKDL DEESKKAYELVLEMLEMVENGVPFEEIYKKYEEMKK |
| AFP1 | SKRSFIEDLLFNKDIDGKYKVIYKYFPEWKPYCDEAKEFVSKLYDELKKKYPNNKEFLKKLDE TALICYLYYVFYLKENPNDKETPKKFCEEKFKLTELS |
| AFP3 | SKRSFIEDLLFNKEFQEILNKIKEAAEKCLKEGPTPEAIKEFEEVVKKCKEELEKFLKDNPEYTE EQKKFIKELFEKHVEECKKKIEEKRRALLEAAGLA |
| AFP5 | SKRSFIEDLLFNKTISEEVLEEIIEVIEKMSDEELLAAFKVIKEAIEKGNPPSPEAFEEAKKKQKT EEGKKLVELARYLLENYPEAFEDFVEGIDEILAS |
| AFP8 | SKRSFIEDLLFNKEKKEEFLKEYKEIKEKVEKEIEEFKKEEEKEAEKKKKEIEKTIKDEKEKEKKI KEVEEEKKEKIEKKKKELEEKLKKELEKLFEKFK |
| AFP9 | SKRSFIEDLLFNKPYRRLVIFLTGMSKLAGNDPAWKALKKKFLETDIKPEDVFDFMLEATRYLL ANTDGTASPEDILAVFRKATEGDPRTDAAVAELRAA |
| AFP10 | SKRSFIEDLLFNKEKLKEYLEKKEKELEKEKEEAKKKLEEILEKHYDKEKNIKYPMSEEEKKKK EEYEKKLKEIEKKLKEVKELKEKLEKEEKEKKELEK |
| AFP11 | SKRSFIEDLLFNKASYREEGDKLIFKSGSVEIIMNKKDNKVVDVKIDPAYKDSPETAKDFRRCL EMIKYFNLPIIFPPEIIAKLPPACQKVLAEFLKELA |

**Table S1. Amino acid sequences of epitope scaffolds.**
